# Structural and proteomic changes in viable but non-culturable *Vibrio cholerae*

**DOI:** 10.1101/433326

**Authors:** Susanne Brenzinger, Lizah T. van der Aart, Gilles P. van Wezel, Jean-Marie Lacroix, Timo Glatter, Ariane Briegel

## Abstract

Aquatic environments are reservoirs of the human pathogen *Vibrio cholerae* O1, which causes the acute diarrheal disease cholera. Upon low temperature or limited nutrient availability, the cells enter a viable but non-culturable (VBNC) state. Characteristic of this state are an altered morphology, low metabolic activity and lack of growth under standard laboratory conditions. Here, for the first time, the cellular ultrastructure of *V. cholerae* VBNC cells raised in natural waters was investigated using electron cryo-tomography complemented by comparison of the proteomes and the peptidoglycan composition of LB overnight culture and VBNC cells. The extensive remodeling of the VBNC cells was most obvious in the passive dehiscence of the cell envelope, resulting in improper embedment of flagella and pili. Only minor changes of the peptidoglycan and osmoregulated periplasmic glucans were observed. Active changes in VBNC cells included the production of cluster I chemosensory arrays and change of abundance of cluster II array proteins. Components involved in iron acquisition and storage, peptide import and arginine biosynthesis were overrepresented in VBNC cells, while enzymes of the central carbon metabolism were found at lower levels. Finally, several pathogenicity factors of *V. cholerae* were less abundant in the VBNC state, potentially limiting their infectious potential.

## Introduction

Changes in the physical and chemical properties of their environment, such as heat, cold and salt stress, oxygen and nutrient deprivation, desiccation and changes in osmolarity, threaten the survival of bacteria, and they have therefore evolved various strategies to evade detrimental effects [1–3]. The facultative human pathogen *Vibrio cholerae*, like many other bacteria, enters a state of restrained metabolic activity when confronted with cues like low temperatures and or low nutrient availability over extended periods of time [4–11]. Once the cells have entered this state, *V. cholerae* does not readily start to grow and reproduce when they are returned to more favorable and nutrient-rich media, and their status has, therefore, been termed “viable but non-culturable” (VBNC) [4, 12, 13].

Under standard laboratory conditions, *V. cholerae* cells are slightly bent, comma-shaped rods. A characteristic morphological feature of *V. cholerae* VBNC cells is a smaller size and a round, coccoid shape with an increased gap between the cytoplasmic and outer membrane (OM), which was previously observed in transmission electron microscopy (TEM) studies [14–16]. It is currently unknown how this reflected in altered peptidoglycan architecture, changes in the membrane structures, or the composition of the periplasmic content of *V. cholerae*. An additional limitation lays in the published TEM images themselves, which were generated using fixed, stained and sectioned cells. This technique is known to affect or obscure the delicate ultrastructure of cells. For example, *Escherichia coli* VBNC cells that were prepared that way were interpreted as dead, due to their seemingly empty cytosol [16]. The presence of several macromolecular complexes in VBNC cells was revealed using other techniques. For example, the toxin co-regulated pilus (TCP) and the flagellum were detected in cultures that are transitioning into, or have already entered, the VBNC state using transcription based methods or immunolabeling [15, 17, 18]. Unfortunately, these methods can only detect the presence of macromolecular complex components, but whether those machineries are still properly assembled and embedded in the envelope remained unclear. It is also unknown if and how these structural changes are regulated. Several studies have focused on either individual regulatory and structural components involved in VBNC formation, or analyzed the transcriptional profile of VBNC cells (see review [10]). Unfortunately, the transcriptional studies are difficult to compare with each other as gene targets were not featured in all data sets or the results were contradicting each other [17–19].

To gain detailed insights into the structural makeup of *V. cholerae* VBNC cells, we studied the morphology and structural adaptation of *V. cholerae* VBNC cells preserved in a near-native state, at macromolecular resolution, and in three dimensions using electron cryo-tomography (ECT). This approach was paired with a proteomic analysis of the VBNC cells, as well as a biochemical analysis of the peptidoglycan composition and the presence and abundance of osmoregulated periplasmic glucans. Together, these complementary analysis methods reveal several characteristic changes of the cell morphology and function of VBNC cells.

## Results

### V. cholerae enters the VBNC state after prolonged incubation

In order to simulate the aquatic habitats as closely as possible in our laboratory setting, we used water samples collected from environmental sources, including fresh water (NFW), brackish water (NBW) and salt water (NSW). Cells introduced into the 50 ml microcosms of NFW, NBW and NSW had all lost their ability to form colonies on LB plates over the course of 200 days (Fig. 1). At the final time point 69% (NFW), 70% (NBW) and 65% (NSW) of the cells were viable according to live/ dead staining with Syto9 and propidium iodide (data not shown).

**Fig. 1:**
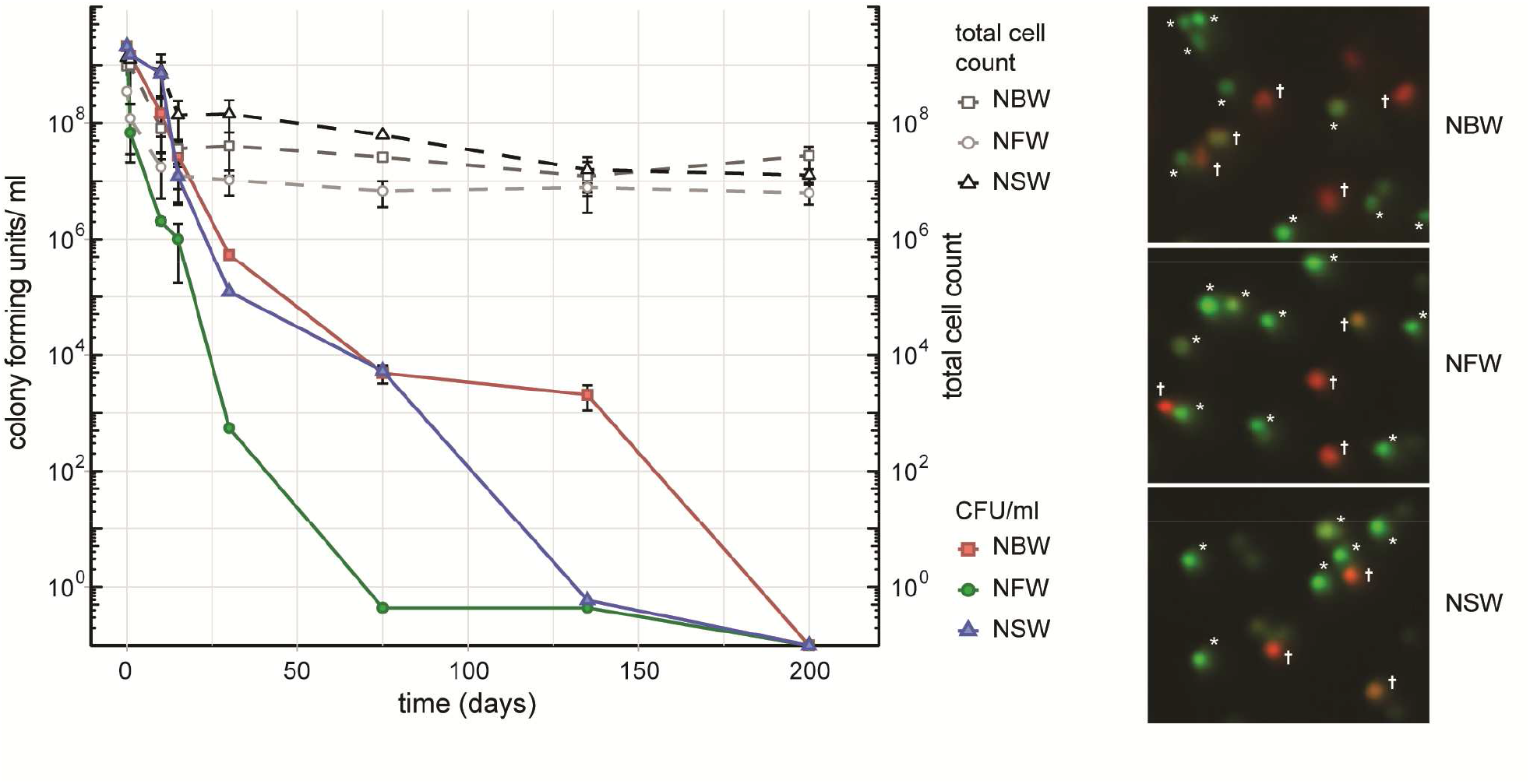
Cell number, CFU and viability during VBNC formation. Left panel shows the colony forming units (CFU)/ ml and total cells count of V. cholerae incubated in natural brackish (NBW), fresh (NFW) or salt water (NSW). Right panel shows examples of the live/ dead staining using propidium iodide and Syto9. Cells marked with an asterisk were counted as “live” while cells with yellow, orange or red hue (marked with a cross) are deemed “dead”.

### Biochemical analysis of OPGs and PG

The transfer of the LB ON cultures to the respective water samples causes an osmotic up- or downshift in our cell cultures. Therefore, we analyzed the presence and quantity of osmoregulated periplasmic glucans (OPGs), which are a known response to changes of environmental osmolarity in growing cells [20]. *V. cholerae* from LB ON cultures used to inoculate the natural water microcosms contained 0.37 µg OPGs/ mg cell weight, while cells grown in LB with no NaCl contained 1.13 µg OPGs/ mg, and cells from LB +30 g NaCl/ l contained 0.20 µg OPGs/ mg cells, respectively. The VBNC cells were all within the range of the value of the inoculum with 0.27 µg OPGs/ mg cells (NFW), 0.17 µg/ mg cells (NBW) and 0.25 µg/ mg cells (NSW) indicating either a lack of OPG production or immediate cession of growth.

The change from ‘comma’-shaped to small, round cells suggests that the PG may be remodeled between these two states. To test this hypothesis, we analyzed digested murein sacculi by LC-MS (Table S1 and Fig. S2). Of monomeric muropeptides, the amount of tetrapeptides increased from 70.3% in stationary cells to 79.4% in VBNC cells. Similarly, the amount of TetraTetra dimers increased from 38.3% in stationary cells to 62.3% in VBNC cells. The PG of both cell types contains low amounts of D-methionine (10% in stationary cells and 6% in VBNC). The amount of muropeptides carrying an anhydro-group at MurNAc was similar between the two cell types, indicating that the length of the glycan strands was generally similar. No pentapeptides were detected in the VBNC cells, which corresponds well to the observed lack of growth. Furthermore, comparison of the base peaks of disaccharides versus bi-disaccharides of both sets, showed that LB ON cells carried 20% more cross-links than the VBNC cells.

### General morphological and physiological changes

Electron cryo-tomography of *V. cholerae* VBNC cells revealed their characteristic morphology; these small, round cells contain a cytoplasmic compartment of reduced volume and are devoid of storage granules, independent of the water used for incubation (Fig. 2A, S1 and movies 1–4). Of the observed cells, 14% (NBW), 20% (NSW) and 22% (NFW) had a damaged cytoplasmic membrane (CM), resulting in loss of cytosolic content (for examples, see Fig. S1). These cells were deemed ‘dead’ and were excluded from the subsequent analysis.

**Figure 2:**
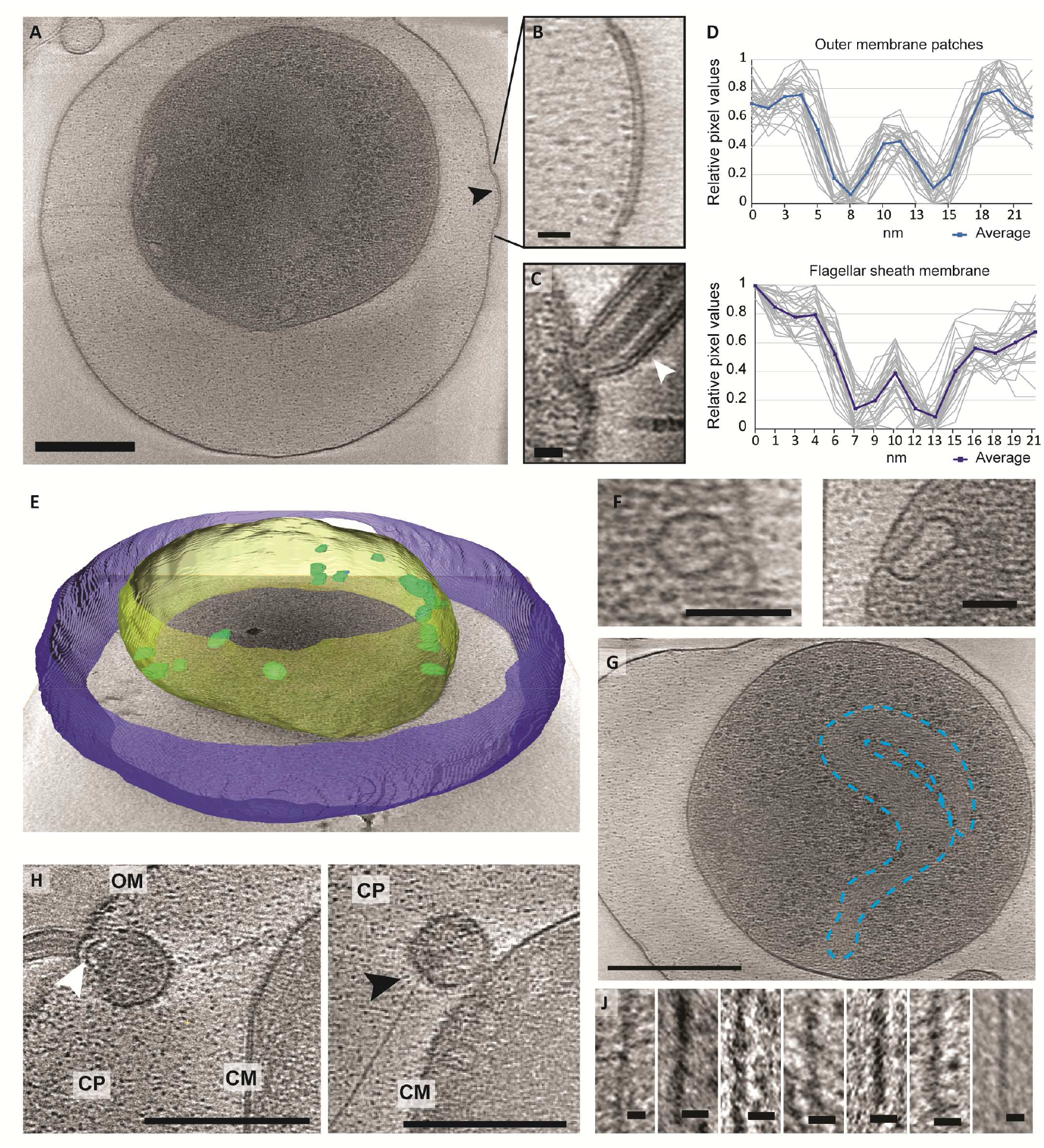
Structural changes of VBNC cells as observed by ECT. A) A cell from a NBW microcosm with a convex OM patch (black arrow). B) Magnified detail of A exhibiting the pronounced two layers of the OM patch. C) Flagellar sheath membrane (white arrow) for comparison. Individual (gray) and averaged (blue) line scan measurements perpendicular to the membrane exhibit similar spacing of the two layers in the outer membrane patch (upper panel) and flagellar sheath membrane (lower panel). E) Tomogram of a NBW VBNC cell segmented using Amira. Outer membrane (blue) is clearly separated from the cytoplasmic membrane (yellow). Multiple invaginations (green) can be found lining the cytoplasmic membrane. F) Examples of the invaginations found in VBNC cells from NBW (left panel) and NSW (right panel) microcosms. G) Tentative outline highlighting the condensed chromosomal DNA. H) VBNC cells showing flagella (left panel, white arrow) or T4P (right panel, black arrow) embedded in small vesicles. J) Details of exemplary T4P. Scale bars represent 200 nm (A and G), 100 nm (H), 50 nm (F) and 10 nm (B, C and J), respectively. CM = cytoplasmic membrane, CP = cytoplasm, OM = outer membrane.

In 29% of all VBNC cells, we observed up to five convex patches in the OM, containing two pronounced layers that differ from the typical OM of *V. cholerae* (Fig. 2A & B). These were never observed in standard overnight LB (LB ON) lab culture cells. Using line scans perpendicular to the membranes, the thickness and spacing between the two layers was determined. We found that the OM patches closely resemble the appearance of the flagellar sheath membrane (Fig. 2C & D).

A characteristic feature of the CM was the presence of multiple invaginations into the cytoplasm that were at least partially open to the periplasm (Fig. 2E&F). These invaginations were found in VBNC cells from NBW and NSW microcosms as well as in cells raised in overnight LB cultures, but not in cells incubated in NFW.

Multiple large proteinaceous structures required for important cellular behaviors such as sensing, motility, attachment and secretion are embedded in the cellular envelope. Since an expansion of the periplasmic space is characteristic for VBNC cells, the functionality or synthesis of these envelope embedded complexes may be affected in the VBNC. Thus, we next focused on determining if they are still present and also if they retained their subcellular location (Table 1). ECT analysis revealed that many VBNC cells were flagellated. In 18% of all VBNC cells, the flagellar basal body or a pilus is contained in a small vesicle and, thus, separated from the bulk cytoplasmic compartment (Fig. 2H). The type IV pili (T4P) that we observed in VBNC cells all had a diameter of ∼6 nm (Fig. 2J) and are therefore unlikely to be TCP pili, which were previously determined to be ∼8 nm thick [21].

Of the three chemotaxis systems present in the *V. cholerae* genome (cluster I, II and III), only cluster II chemosensory systems were found with similar abundance in cells of all incubation types. These can be clearly identified by the presence of periplasmic receptor domains, as well as the characteristic 25 nm spacing between the base plate and the inner membrane [22]. The cytoplasmic cluster I chemosensory system was only observed in a few cells incubated in NBW and NFW, while cluster III was not seen in any of the imaged cells.

Finally, we also observed a region characterized by a relatively homogeneous density and lack of ribosomes, which appears to be the considerably condensed chromosome (Fig. 2G). Intriguingly, we only observed this in the cells from NSW and NBW incubations, but did not observe a similar DNA condensation in the VBNC cells from the NFW sample.

**Table 1:** Quantification of occurrence of structures in cells from VBNC microcosms and LB ON cultures as observed by ECT.

| Sample | N | flagella | chemosensory system |  |  | T4P | detached | condensed |
| --- | --- | --- | --- | --- | --- | --- | --- | --- |
|  |  |  | I | II | III |  | OM | DNA |
| NBW | 35 | 66% | 9% | 71% | - | 23% | 100% | 83% |
| NFW | 18 | 33% | 6% | 94% | - | 72% | 100% | - |
| NSW | 18 | 39% | - | 72% | - | 22% | 100% | 83% |
| LB ON | 18 | 72% | - | - | - | 33% | 17% | - |

### Proteomic analysis of VBNC cells

To gain more insights into changes associated with the VBNC state, the proteomes of VBNC cells and cells of an LB ON lab culture were compared.

Since many of the structural changes were highly similar in VBNC cells obtained from all types of water samples, NBW water cultures were used for proteomic analysis using mass spectrometry and label-free quantification. The respective names were inferred from Uniprot, and functional categories were assigned using KEGG and DAVID [23–26]. Of the 2219 quantified proteins, 1349 were significantly over- or underrepresented (q<0.01) in VBNC cells as compared to LB ON cells, demonstrating the extensive impact of the VBNC status on the cellular content (Tables S2 and S3, Fig. S3). Proteins that were detected with n>=3 peptides were analyzed in more detail. Components that were more abundant in VBNC cells fell, among others, into the categories of bacterial chemotaxis, flagellar assembly, ribosomal proteins, arginine biosynthesis and ABC transporters. Less abundant proteins in VBNC cells were predominantly allocated to metabolic categories such as carbon metabolism, TCA cycle, amino acid, amino sugar and nucleotide sugar metabolism (Table S4). For more detailed analysis, the components of individual structures or pathways were further refined using published data or the VchoCyc database [27].

#### Large protein complexes

In the proteomic analysis, we detected multiple proteins involved in the formation of T4P, flagellar and chemosensory systems with significant changes between LB ON cultures and VBNC cells (Fig. 3).

The *V. cholerae* genome contains three T4P systems: the mannose-sensitive haemagglutinin (MSHA) pili, the toxin co-regulated pili (TCP) and the chitin co-regulated pili (ChiRP) [28]. The seven detected MSHA pili proteins presented only a slightly lower or higher concentration in VBNC compared to LB ON cells, or were not significantly different between the cultures. Very few proteins of the other two pili systems were detected. Of those, TcpF levels in VBNC cells was not significantly different from LB ON cultures, but all other TCP and ChiRP components were less abundant.

*V. cholerae* moves through the environment using flagellar rotation guided by chemotaxis. Seven out of 19 detected flagellar proteins showed a > 1.5 – fold abundance, further 10 were represented at similar levels as in LB ON cells and only FlhF and FlhA were significantly less abundant in VBNC cells. Flagellar motility is controlled by the chemosensory cluster II. Five of the 40H class methyl-accepting chemosensory proteins (MCPs) associated with cluster II were > 1.5 – fold more abundant while three were 1.5-fold reduced. The remaining 22 detected cluster II components, including CheA and CheW, showed only minor changes. As observed in the ECT data, a fraction of the VBNC cells from NBW microcosms contained cluster I chemosensory arrays which is in line with the increased abundance of the cluster I chemosensory array CheA (VC1397), CheW (VC1402) and the MCP DosM (VC1403) in these cells.

#### Pathogenicity

Besides the TCP pilus, a plethora of other virulence factors and regulators are known or likely to be involved in pathogenicity of *V. cholerae*. This includes the cholera toxin CtxAB, the T2SS EpsCDEFGHIJKLMN, hemolysin A, ten further predicted hemolysins or hemolysin secretion proteins, the accessory colonization factor AcfBC, the MARTX, Zot and Ace toxins. Only 15 of these were detected with more than two peptides in the MS analysis. Of these, only the putative hemolysin secretion protein VC0199, a putative hemolysin (VC0578), the hemolysin Hlx VCA0594 and the T2SS proteins EpsCEFGL were found with a slightly increased abundance in VBNC cells (Fig. 4).

**Figure 3:**
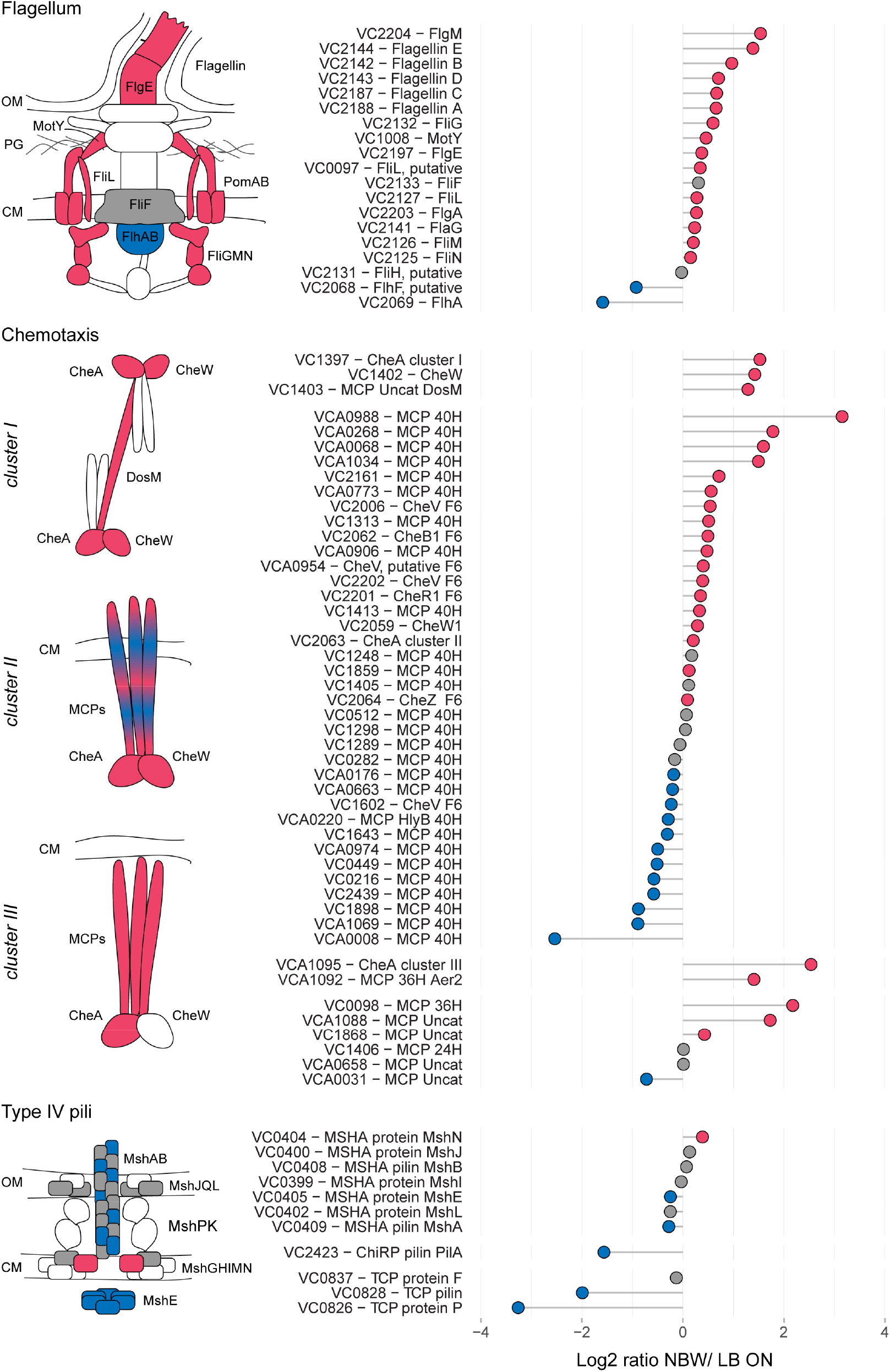
Relative protein abundances of components of the flagellum, chemotaxis arrays and T4P. The left illustrations depict which components of the respective structures are more (red) or less (blue) abundant or not significantly different (grey). White symbols represent components that were not found in the MS analysis or with peptides n<2. CM = cytoplasmic membrane, OM = outer membrane. Depicted is the log_2_-ratio of NBW VBNC cells and LB ON cells. Proteins found in significantly higher abundance in VBNC cells compared to LB ON are indicated by red circles while significantly lower ratios are shown as blue circles. Grey circles represent components with no significant difference between VBNC and LB ON cells (q<0.01). Chemotaxis components are grouped by their predicted interaction with the respective chemosensory cluster.

**Figure 4:**
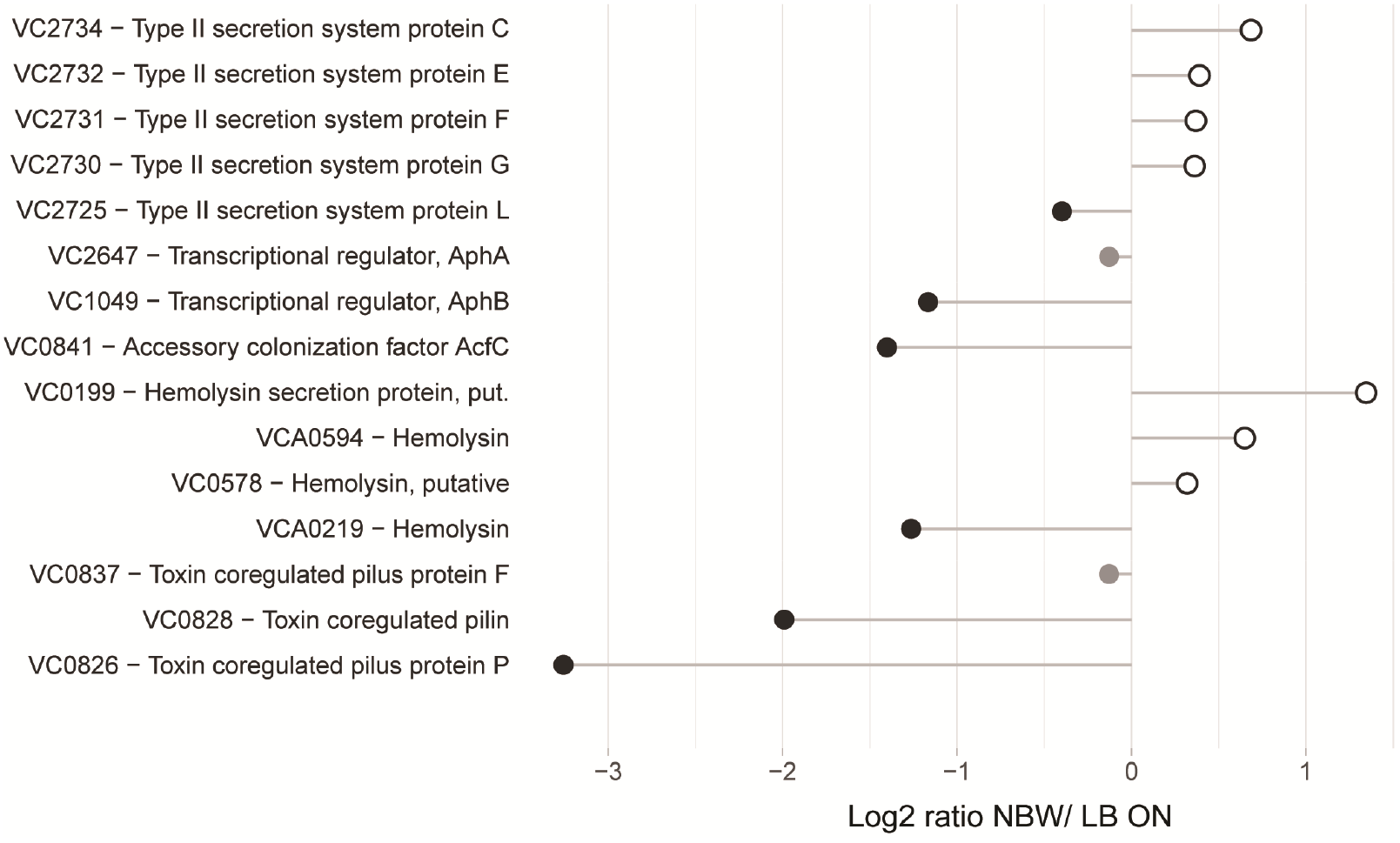
Relative protein abundances of proteins associated with pathogenicity. Depicted is the log_2_-ratio of NBW VBNC cells and LB ON cells. Proteins found in significantly higher abundance in VBNC cells compared to LB ON are indicated by open white circles while significantly lower ratios are shown as black circles. Protein abundances that were not significantly different are colored grey (q<0.01).

#### Envelope maintenance, cell shape and division

Contrary to the drastic morphological changes of the envelope of VBNC cells, half of the proteins involved in cell division, cell shape and the synthesis and maintenance or structure of the cell envelope were not significantly more or less abundant (Fig. S5). If we observed differences, these changes differed only slightly between the cultures (up to ±1.5 fold). This includes the proteins that determine the rod-shape (MreBC) and curvature of the cell (CvrA). The two enzymes predicted to be responsible for OPG production, MdoHG (VC1287 and VC1288) were less abundant than in cells grown in LB overnight cultures.

#### Metabolic enzymes

VBNC cells have previously been reported to exhibit a markedly restrained metabolism [5–7, 11]. To analyze potential shifts in metabolic pathways, we retrieved the predicted function of metabolic proteins detected in the VBNC proteome data, mapped their respective pathways and expected underlying regulons using KEGG, VchoCyc and RegPrecise 3.0 [25, 27, 29, 30] (Table S5 and S6). Cell envelope proteins involved in electron transport such as cytochromes (VC1442, VC1441, VC1440, VC1439 & VC1951), formate dehydrogenases (VC1519, VC1511 & VC1512) and cytochrome d ubiquinol oxidases (VC1843 & VCA0872) were less abundant in VBNC cells. We observed a significant reduction of enzymes involved in the central carbohydrate metabolism, specifically those required for glycolysis, the TCA cycle and the pentose phosphate pathway. Furthermore, we found a reduction of proteins predicted to be involved in the biosynthesis of lipopolysaccharides, isoprenoids, nucleotides and branched-chain amino acids as well as of co-factors such as riboflavin, NAD, ubiquinone and glutathione. In contrast, proteins of the Entner-Doudoroff (ED) pathway, the urea cycle and fatty acid β-oxidation were more abundant in VBNC cells. In addition, components of the amino acid metabolism, as well as the siderophore and iron-sulfur cluster biosynthesis, were increased. Predicted ABC transporters with at least two components significantly more abundant, were those involved in up-take of sulfate, iron (III), arginine/ornithine, L-amino acids and oligopeptides (Table S7).

Since the components of the mentioned pathways are often part of the same regulon(s), we next analyzed their abundance in relation to the regulatory networks. Genes that are upregulated in *V. cholerae* O395 by low iron availability and/ or the absence of master regulator Fur, or contained a predicted Fur binding box in their upstream region, were used to define the subclass of iron/Fur regulated proteins (Fig. 5) [31, 32]. With the exception of a ferroxidase (VC0365) and a protein putatively involved in iron uptake (VC1264), all other proteins whose genes are repressed by Fur are present in higher concentrations in VBNC. This also includes the gene for manganese-dependent superoxide dismutase SodA, while the iron-dependent superoxide dismutase SodB was two-fold less abundant. Proteins encoded by genes that are either activated by Fur or not part of its regulon (VCA0678, VCA0679, VC1516, VCA0784 and VC1216) were found in lower or only slightly higher abundance in VBNC cells. Proteins encoded by genes with a predicted upstream Fur binding box in their upstream region, but without expression data from a Fur mutant, showed a mixed pattern.

**Figure 5:**
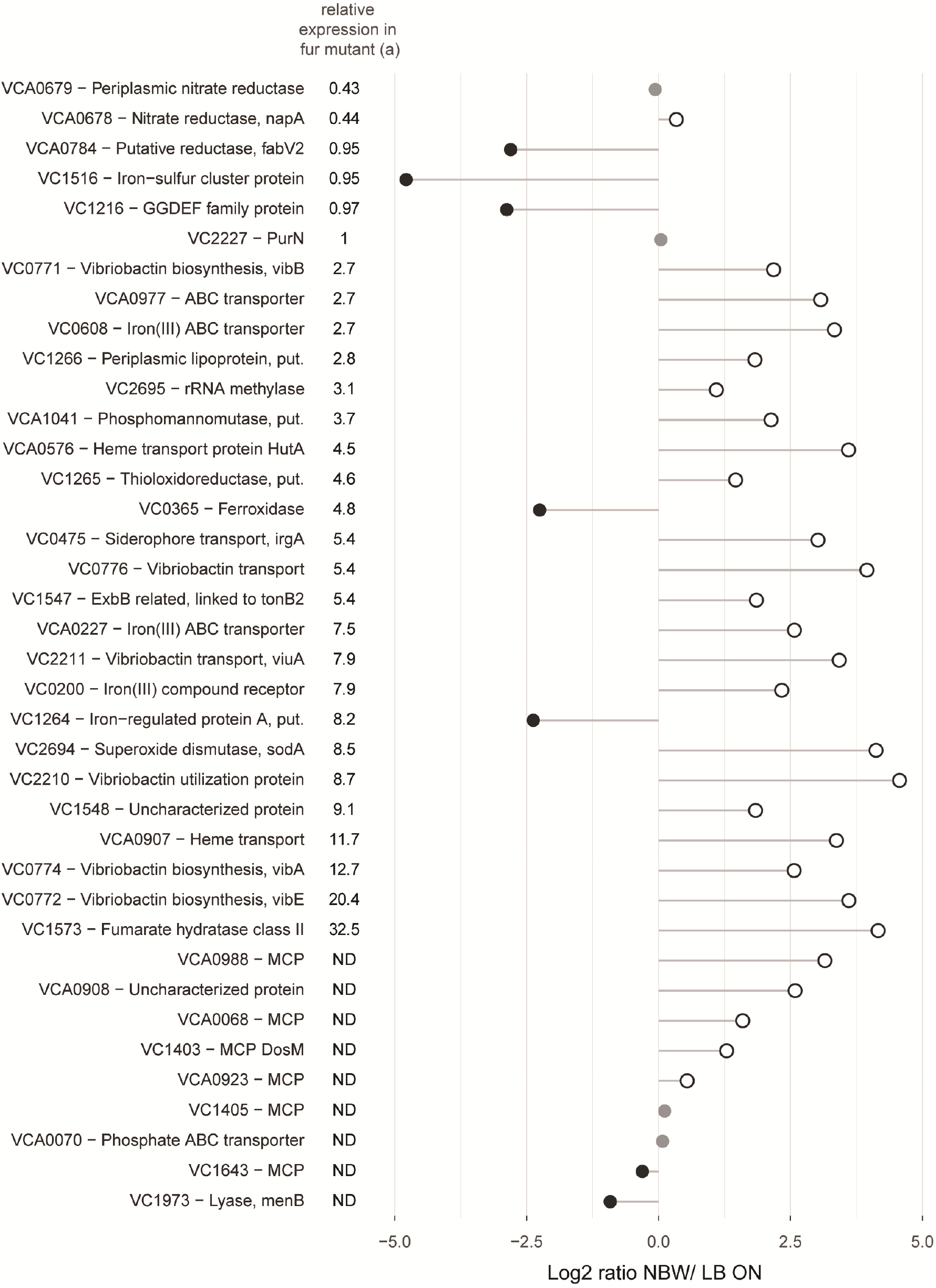
Relative abundances of proteins with predicted Fur binding boxes in their upstream region and/ or shown regulation by iron availability and/ or the master regulator Fur. Depicted is the log_2_-ratio of NBW VBNC cells and LB ON cells. Proteins found in significantly higher abundance in VBNC cells compared to LB ON are indicated by open white circles while significantly lower ratios are shown as black circles. Protein abundances that were not significantly different are colored grey (q<0.01). (a) Relative expression data was taken from Mey et al. [31]. The genes of the first five proteins were positively regulated by iron and independently of Fur while the following 24 components were negatively regulated by Fur and iron. ND = not defined.

Besides the already described Fur regulon, all proteins whose genes are predicted to be repressed by ArgR, GntR, IscR and TyrR, as well as those for the repressors themselves, were more abundant. Genes that are predicted to be repressed by FruR, GalR, NanR, PdhR, PrpR and ScrR showed a decrease in expression, as based on the proteome data. With the exception of NanR, all these repressors were also less abundant. The remaining regulons showed a mixed pattern with varying protein levels of their regulons.

## Discussion

Cells that have reached the VBNC state have been subject to osmolarity up- or downshifts, low temperature and starvation for extended periods of time. As a result, the cells have undergone a process of drastic physiological and morphological restructuring. Three-dimensional structural analysis of cells in a near-native state at macromolecular resolution revealed the vast changes of the cell envelope and macromolecular structures. The cytoplasm of *V. cholerae* VBNC cells was relatively small and devoid of storage granules. The reduced cell size could potentially be created during cytoplasm shrinkage. Here, the flagellum, pili and other membrane spanning structures could act as tethers that anchor the CM to the OM. This, in turn, could lead to the observed second cytoplasmic compartment of smaller size. In general, the abnormal distance between CM and OM may disrupt a variety of processes such as envelope maintenance, sensing and motility as previously shown for *E. coli* [33].

The OM of VBNC cells showed thick, convex patches reminiscent of the flagellar sheath. The asymmetric composition of the OM is tightly regulated to maintain its barrier function (see review [34]). Aberrations, such as the OM patches, might be symptomatic of a perturbed OM maintenance.

The detected proteins required for envelope maintenance and shape exhibited either unchanged or only minor differences in VBNC cells. Thus, the lack of conservation of the OM, CM and PG is likely due to the increased gap of the periplasm, rather than the absence of the maintenance machinery. Contrary to the radical change of the general morphology of *V. cholerae* VBNC, the PG alterations are rather subtle. No pentapeptides, were detected, indicative of lack of cell wall synthesis. The PG appeared less cross-linked in VBNC cells, while the glycan strands had a similar length as in PG obtained from LB ON cells. This is in contrast to the substantially higher rate of cross-linking, as well as shorter glycan strands, that were reported for *E. coli* VBNC cells [35]. The authors speculated that especially the shorter glycan strands contribute to the formation of coccoid cells. In *V. cholerae* VBNC cells, the rounded cell shape may arise due to the lower degree of cross-linking of the PG which reduces cell wall stiffness [36, 37].

The invaginations of the CM found in cells incubated in NBW, NSW and LB ON incubations may be the result of an increase in protein content relative to the lipid membrane. A similar mechanism has previously been reported in cells that overproduce membrane proteins [38–40]. The low level of OPGs in the periplasm of NFW VBNC cells may weaken their osmoprotection and prevent the formation of invaginations as the CM might be more strained through increased turgor pressure. The described cell envelope anomalies may affect the function of the envelope-spanning structures such as pili and the flagellum. Certain integral structures, such as the alignment subcomplex of T4P and the stators of the flagellum, span the distance from the CM to the PG and their proper placement is essential to the respective functions [41–45]. Thus, the pili and flagella that reside either in the small cytoplasmic vesicles or at sites of an enlarged periplasmic gap are likely not functional because of the increased distance between the envelope layers. The pili found in the VBNC cells are likely MSHA pili. We base this conclusion on their measured width of about 6 nm which matches the previously reported diameter of T4a pili [21], as well as the stable presence of several MSHA pili components and decreased abundance of ChiRP and TCP system components in the proteomics analysis.

A previous report by Kim et al. on *E. coli* VBNC populations concluded that VBNC cultures are mainly composed of dead cells, as the majority of cells appeared to have an empty cytosol [16]. This observation solely relied on the analysis of chemically fixed, dehydrated, heavy metal stained, resin-embedded and thin sectioned cell material without the three dimensional context of the whole cells. Each of these sample preparation steps can introduce artifacts and destroy the delicate ultrastructure of bacteria [46, 47]. Our analysis of 3D tomograms of flash-frozen VBNC cells cannot confirm the statement by Kim et al. for *V. cholerae* VBNC cells, since we only observed cells with a damaged CM and cytosol leakage in 14–22% of the cases. This number resembles the dead fraction determined by the live/dead staining experiment.

In addition to the changes in the *V. cholerae* VBNC cell envelope, we found several changes in the protein content of the cytoplasm. *V. cholerae* encodes three distinct chemotaxis systems (cluster I, II, III) and 44–45 different MCPs [48]. The MCPs that are known to interact with cluster II all belong to the 40H class [22, 49]. Cluster II arrays sense a plethora of cues and are so far the only chemosensory system that controls the chemotactic swimming behavior of *V. cholerae* [50]. This array is present in both VBNC and LB ON cells at similar frequency and with comparable CheA levels, but it seems to undergo restructuring based on the differences of the 40H MCP abundance. A recent study of *V. cholerae* cluster II revealed that this array is not ultrastable like in the case of *E. coli* chemotaxis arrays. This may facilitate MCP turn over and allow this species to quickly remodel this system to tune their chemotactic response. As the molecules sensed by individual MCPs are still unknown for most MCPs, the cues that these restructured cluster II arrays can sense remain to be uncovered. Cluster I chemosensory arrays were only observed in VBNC cells where the respective proteins were also 2.4–2.8 times more abundant. These arrays have been reported to be formed under low-oxygen conditions [51]. As our cells were incubated in closed microcosms, they likely experienced oxygen depletion, which we assume induced the cluster I formation. Furthermore, we also observed a homogenous region in the cytoplasm that is reminiscent of the compact nucleoid of other bacteria [47, 52]. Thus, we believe this region to be the condensed chromosomal DNA. Multiple proteins are capable of structuring and condensing DNA. Out of these, the IHF subunits and DPS were significantly more abundant in VBNC cells. Together, these proteins could be responsible for observed compacting the chromosome. However, DNA condensation is thought to be an interplay between molecular crowding and DNA structuring proteins [53]. This might explain why the compacted DNA was not observed in fresh water treated cells.

Similar to the absence of TCP pili, most other known pathogenicity factors were either not detected or less abundant in the proteomic analysis of VBNC cells including the cholerae toxin and hemolysin A. This is in contrast to recent transcriptomic analysis of *V. cholerae* VBNC cells from cold seawater. In this study, the TCP pilus, the cholera toxin, and toxin related genes were found to be expressed at a similar level or even upregulated. Based on these results, the authors concluded that VBNC cells retain their pathogenic potential [15, 18, 54, 55]. To date, a single trial with nine participants determining the infectivity of *V. cholerae* VBNC cells has been published. Only one of the volunteers developed cholera symptoms. However, the authors noted that the loss of pathogenicity may be dependent on how long the cells are already in the VBNC state [56]. A time course dependent determination of the presence of pathogenicity factors is, to date, still lacking.

*V. cholerae* VBNC cells not only restructure their morphology but also their metabolism. Our proteomic data shows that proteins of several entire predicted regulons or pathways exhibit abundance shifts. This indicates the restructuring of the metabolism caused by loss of repression of several regulons. This includes most elevated gene products repressed by Fur, ArgR, GntR, IscR and TyrR. The carbon metabolism also seemed to undergo a broad shift as proteins involved in glucose utilization (PtsGHI) and the Embden–Meyerhof–Parnas-pathway (Pgi, TipA, GapA) were all less abundant, while the glyoxylate shunt components showed stable (AceA) or slightly higher levels (AceB). In addition, VC0285, a key protein of the ED pathway, and several enzymes of FA degradation were detected with increased levels. Furthermore, several ABC transport systems implicated to take up amino acids or peptides were found at higher levels. This indicates a vast metabolic shift towards alternative energy sources. This seems logical as *V. cholerae* was shown to lose 88.7% of their carbohydrates within a few days of starvation [57].

A general question regarding VBNC formation is whether the observed changes are due to active regulation or passive mechanisms such as stalled synthesis or maintenance and stochastic decay. Our data suggests that many of the alterations observed in the *V. cholerae* VBNC cell envelope are not due to active restructuring, but rather to passive changes that may be due to the dehiscence of the two membranes caused by a shrinking cytoplasm. The metabolic changes, induction of the cluster I chemosensory system and restructuring of the cluster II array appear actively regulated and in dependence to effector and oxygen availability. Whether the lower abundance of toxins and the absence of the TCP pilus is an active downregulation cannot be assessed, but indicates that the infectious potential of *V. cholerae* VBNC cells is likely diminished.

Most previous work consists of individual aspects of VBNC formation and decoupled structural and proteomic or transcriptomic studies. Here, we have coupled the proteomic and structural analysis to give important new insights into the extensive changes this pathogen undergoes upon VBNC-inducing conditions. To further study the temporal progress of VBNC development, future studies should include the proteomic analysis along the entire time course of their formation.

## Material and Methods

### Bacterial strain, media and VBNC inducing conditions

*Vibrio cholerae* O1 biovar El Tor str. N16961 was used in this study. A small amount of these cells from LB plates was used to inoculate three overnight cultures of 250 ml low salt LB. These cultures were grown at 30°C for 18h shaking at 200 rpm with a final OD_600_ of 2.5, corresponding to ∼2·10^9^CFU. 50 ml of the cells were harvested in 50 ml conical polypropylene tubes using centrifugation (20 min at 5000xg). Cell pellets were carefully resuspended in cold natural waters, closed and kept at 4°C in the dark on a rocking platform set to low agitation. Natural waters were collected at the following sites: NFW at 51°52’50.3”N 5°59’52.2”E, NBW containing 13 g chloride/ l was collected at 53°06’02.4”N 4°53’50.5”E and NSW at 53°06’11.7”N 4°53’54.8”E. All waters were filtered through a series of whatman filters, sterile filtered using a 0.22µm filter followed by autoclaving. Waters were stored at 4°C until use.

### Live/dead stain

The fraction of live cells was determined as previously described [58]. In essence 200 µl of a 1:50 dilution of each incubation was mixes with 1 µl of Syto9 and propidium iodide each. The mixture was incubated at RT for 20 min in the dark before 3 µl were applied to an agarose slab and imaged on a Zeiss Axioplan 2 equipped with a Zeiss AxioCam MRc 5 digital color camera. At least 500 cells from randomly chosen areas were quantified. Images were processed using ImageJ. Cells with a red, orange or yellow hue were rated as “dead” and green cells as “live”.

### Electron cryo-tomography

18 µl of cell suspension from the respective cultures was gently mixed with 2 µl protein A-treated 10 nm colloidal gold solution (Cell Microscopy Core, Utrecht, The Netherlands) by pipetting. Aliquots of 3 µl were applied to a plasma-cleaned R2/2 copper Quantifoil grid (Quantifoil Micro Tools, Jena, Germany). Plunge freezing into liquid ethane was carried out using a Leica EMGP (Leica microsystems, Wetzlar, Germany) set to 1 s blotting inside the chamber set at 20°C and 95% humidity. Grids were stored in liquid nitrogen until imaging.

Data acquisition was performed on a Titan Krios transmission electron microscope (Thermo Fisher Scientific, Hillsboro, OR, USA) operating at 300 kV. Images were recorded with a Gatan K2 Summit direct electron detector (Gatan, Pleasanton, CA) equipped with a GIF-quantum energy filter (Gatan) operating with a slit width of 20eV. Tomograms were recorded at a nominal magnification of 42,000x (pixel size of 3.513 Å). Using UCSF tomography data collection software [59], all tilt series were collected using a bidirectional tilt scheme which started with 0° to −54° followed by 0° to 54° tilting with a 2° increment. Defocus was set to –8 µm. The cumulative exposure was 120 e-/ Å^2^. Drift correction and bead-tracking based tilt series alignment were done using software package IMOD [60]. Tomograms were reconstructed using simultaneous iterative reconstruction (SIRT) with iteration number set to 4.

Line scans perpendicular to the OM or sheath membrane was done using IMOD. 29 measurements from five cells each were recorded.

### OPG analysis

Bacteria (100 mL) were grown in LB with several NaCl concentrations or in various natural waters. Bacteria were collected by centrifugation at 4°C for 15 min at 8,000 g. Cell pellets were suspended in 20 mL of distilled water and lysed with 5% trichloroacetic acid (TCA). After centrifugation at 4°C for 30 min at 8,000 g, the supernatant was neutralized with ammonium hydroxide 10% and concentrated by rotary evaporation. The resulting material (2 mL) was fractionated by gel filtration on a Bio-Gel P-4 column (length: 47 cm, diameter: 1.7 cm) equilibrated with 0.5% acetic acid. The column was eluted in the same buffer at a flow rate of 15 mL h-1 and fractions of 1.5 mL were collected. Presence of oligosaccharides in each fraction was determined by the colorimetrically anthrone procedure [61]. Fractions containing OPGs were pooled and total content was determined by the same procedure.

### Peptidoglycan analysis

Murein sacculi were purified as described previously [62]. In essence, 50 ml cell material of LB overnight cultures or VBNC (NSW) cell suspensions was pelleted, resuspended in 5 ml PBS and slowly added to 10 ml of boiling 10% SDS while stirring. Samples were boiled for 4 h, then stirred overnight at 37°C. Cell wall material was then pelleted by ultracentrifugation (85.000 rpm, 0.5 h) and washed in MQ water. Sacculi were digested with pronase E (0.1 mg/ml) in a Tris-HCl 10 mM pH 7.5 buffer for 1 hour at 60°C to remove Braun’s lipoprotein. After heat-inactivation and washing, the samples were treated with muramidase (100 mg/ml) for 16 hours at 37°C, in 50 mM phosphate buffer, pH 4.9. Muramidase digestion was boiled and centrifuged and the supernatants were reduced with 0.5 M sodium borate pH 9.5 and sodium borohydride. Finally, samples were adjusted to pH 3.5 with phosphoric acid.

Chromatographic separation was performed as previously described [63] on an Acquity UPLC HSS T3 C18 column (1.8 µm, 100 Å, 2.1 × 100 mm).

The peak areas of masses corresponding to muropeptides were collected and a final table which shows peak areas as percentage of the whole was produced in Microsoft Excel.

### Proteomic analysis

Equal amounts of *V. cholerae* O1 biovar El Tor str. N16961 cells from LB overnight or VBNC microcosms were collected using centrifugation. Cells were carefully washed with ice-cold PBS and subjected to proteomic analysis using mass spectrometry (MS). In the first step, cell pellets were reconstituted in 2% Sodiumlauroylsarcosinate and heated for 15 min at 95°C. Protein concentration was measured and 50 μg of total solubilized protein was used for further analysis. The samples were incubated with 5 mM tris(2-carboxyethyl)phosphine at 95°C for 15 min followed by 30 min incubation with 10 mM iodoacetamide at 25°C. For in-solution digestion (ISD), the samples were diluted to 0.5% detergent using 100 mM NH4HCO3. 1 µg trypsin (Promega) was used for digestion overnight at 30°C. Prior to Liquid Chromatography-Mass Spectrometry (LC-MS) analysis, samples were acidified and purified using C18 microspin columns (Harvard Apparatus) according to the manufacturer’s instructions.

LC-MS/MS analysis of protein digests was performed on Q-Exactive Plus mass spectrometer connected to an electrospray ion source (Thermo Fisher Scientific). Peptide separation was carried out using Ultimate 3000 nanoLC-system (Thermo Fisher Scientific), equipped with packed in-house C18 resin column (Magic C18 AQ 2.4 µm, Dr. Maisch). The peptides were first loaded onto a C18 precolumn (preconcentration set-up) and then eluted in backflush mode with a gradient from 98 % solvent A (0.15 % formic acid) and 2 % solvent B (99.85 % acetonitrile, 0.15 % formic acid) to 25 % solvent B over 105 min, continued from 25 % to 35 % of solvent B up to 135 min. The flow rate was set to 300 nL/ min. The data acquisition mode for the initial LFQ study was set to obtain one high-resolution MS scan at a resolution of 60000 (*m/z* 200) with scanning range from 375 to 1500 *m/z* followed by MS/MS scans of the 10 most intense ions. To increase the efficiency of MS/MS shots, the charged state screening modus was adjusted to exclude unassigned and singly charged ions. The dynamic exclusion duration was set to 30 sec. The ion accumulation time was set to 50 ms (both MS and MS/MS). The automatic gain control (AGC) was set to 3 × 10^6^ for MS survey scans and 1 × 10^5^ for MS/MS scans.

For label-free quantification the MS raw data were analyzed with Progenesis QI software (Nonlinear Dynamics, version 2.0). MS/MS search of aligned LC-MS runs was performed using MASCOT against a decoy database of the uniprot *Vibrio cholerae* protein database The following search parameters were used: full tryptic specificity required (cleavage after lysine or arginine residues); two missed cleavages allowed; carbamidomethylation (C) set as a fixed modification; and oxidation (M) set as a variable modification. The mass tolerance was set to 10 ppm for precursor ions and 0.02 Da for fragment ions for high energy-collision dissociation (HCD). Results from the database search were imported back to Progenesis, mapping peptide identifications to MS1 features. The peak heights of all MS1 features annotated with the same peptide sequence were summed, and protein abundance was calculated per LC-MS run. Next, the data obtained from Progenesis were evaluated using SafeQuant R-package version 2.2.2 [64]. Hereby, 1% FDR of identification and quantification as well as intensity-based absolute quantification (iBAQ) values were calculated.

## Acknowledgements

The authors acknowledge the support and the use of resources of Instruct-ERIC. Data acquisition was performed at the Netherlands Centre for Electron Nanoscopy in Leiden (NeCEN). This work is part of the research programme “Building Blocks of Life” with project number 737.016.004, which is (partly) financed by the Netherlands Organisation for Scientific Research (NWO). SB was supported by a postdoctoral fellowship from the German Academy of Sciences Leopoldina.

